# Analysis of co-regulated abundance of genes associated with arsenic and phosphate metabolism in Andean Microbial Ecosystems

**DOI:** 10.1101/870428

**Authors:** L.A. Saona, S. Valenzuela-Diaz, D. Kurth, M. Contreras, C. Meneses, E. Castro-Nallar, M.E. Farías

**Affiliations:** Laboratorio de Investigaciones Microbiológicas de Lagunas Andinas (LIMLA), Planta Piloto de Procesos Industriales Microbiológicos (PROIMI), CCT, CONICET, San Miguel de Tucumán, Tucumán, Argentina; Facultad de Ciencias de la Vida, Center for Bioinformatics and Integrative Biology, Universidad Andrés Bello, Santiago, Chile; Centro de Ecología Aplicada, Santiago, Chile; Facultad de Ciencias de la Vida, Centro de Biotecnología Vegetal, Universidad Andres Bello, Avenida República 330, 8370035 Santiago, RM, Chile; FONDAP Center for Genome Regulation, Santiago, Chile

**Keywords:** arsenate and phosphate, Pst transport system, arsenic respiration, Andean Microbial Ecosystem

## Abstract

Phosphate and arsenate are very similar compounds, and there is great interest in studying their relationship and their interaction with biological systems. Despite having no apparent biological function, specific genes regulate arsenic interaction with cells and can be located in regions of the genome called arsenic islands, where phosphate metabolism genes are also present. Although they are neighboring genes, the nature of their relationship and how they have been selected is still unknown.

In this work, we analyzed the metagenomes of the four microbial ecosystems inhabiting hypersaline lakes of the Argentine Puna and the Atacama salt flat in Chile and have evaluated the presence and abundance of both arsenic and phosphate metabolism genes. The samples analyzed included microbialites, biofilms and microbial mats; all of them established under high arsenic concentrations, high UV radiation and high temperature fluctuation, among others.

The results show great differences in the dispersion and abundance of genes related to both phosphate and arsenic metabolism in the analyzed samples. The main difference is given in the Diamante Lake, located in the crater of the Galan volcano characterized by being one of the lakes with the highest arsenic concentration (2.34 mM). Correlating genes abundance with the physicochemical parameters of the lakes studied, our results suggest that arsenic and phosphate metabolism are intricately co-regulated in environmental conditions.

## Introduction

Phosphorus is typically found in nature as inorganic phosphate (Pi). Together with carbon, hydrogen, nitrogen, oxygen, and sulfur, Pi is one of the six chemical elements essential to life as we know it (Karl 2000; Rasuk et al. 2016). Pi is involved in many cellular functions as it is part of DNA and RNA, ATP and polyphosphates (polyP) (Kornberg 1999), membrane phospholipids and proteins. In addition, the metabolism of both prokaryotes and eukaryotes depends on reactions where Pi is essential, e.g., phosphorylation/dephosphorylation reactions, which regulate protein functions in intracellular signaling pathways.

Arsenic is an element structurally similar to phosphorus, they both share characteristics such as atomic radius, and identical electronegativity and oxidation states. In their most characteristic oxidized form (+5), arsenate [As(V)] as well as phosphate (PO_4_^3-^) are negatively charged at physiological pH. Indeed, both molecules present similar speciation at different pH, as well as similar *pK*_*a*_ values (Wolfe-Simon et al. 2008, 2011; Tawfik and Viola 2011) and thermochemical radii, differing by only 4% (Kish and Viola 1999). Further, inorganic arsenate is able to form key biological ester bonds which are analogous to those formed by Pi, including genetic material. This occurs spontaneously between the 5’-hydroxyl group of ribose sugars and arsenate, giving rise to the formation of the mononucleotides 5’-arsenate (Lagunas et al. 1984; Wolfe-Simon et al. 2008).

In spite of such similarities, arsenic is one of the most toxic elements for most lifeforms (Parke 2013). Biomolecules based on As(V) have a much higher rate of hydrolysis than those based on phosphate (Lagunas et al. 1984; Fekry et al. 2011; Parke 2013), and –unlike phosphate– arsenic presents redox reactions in the physiological range of redox potentials, thus, gradually transforming As(V) into As(III) (Schoepp-Cothenet et al. 2011) in the prokaryotic cytoplasm.

In 2010 a super arsenic-resistant bacterium belonging to the Halomonadaceae family, denominated GFAJ-1, was isolated by Wolfe-Simon (2011). According to the authors, the bacterium –isolated from Mono Lake, which is characterized by high arsenic concentrations (200 µM) (Wolfe-Simon et al. 2011; Elias et al. 2012)– might replace the phosphorus in its DNA with arsenic, enabling it to grow in culture media with arsenate and without the addition of phosphate. That article challenged the role of arsenic in biology, and even the concept of life we had until that moment. Although arsenate as an essential component of DNA is totally discredited (Schoepp-Cothenet et al. 2011; Erb et al. 2012; Reaves et al. 2012; Kim and Rensing 2012), microbial cells exhibit specific cellular mechanisms that modulate the As intracellular concentration. Due to its similarities with phosphate, arsenate can enter cells through phosphate transporters; consequently, various organisms possess enzymes specifically related to arsenic resistance and arsenic metabolism, and the genes that code for these enzymes are distributed along the three domains: Bacteria, Archaea and Eukarya (Jackson and Dugas 2003).

Phosphate uptake occurs mainly by two pathways: through a phosphate specific transport (Pst) system or a phosphate inorganic transport (Pit) system (Willsky and Malamy 1980b, a; Guo et al. 2011). Depending on extracellular phosphate concentrations, one transport system is activated over the other, a regulation process that has been reported in different microorganisms such as Proteobacteria, Cyanobacteria, Algae, and Archaea (Rosenberg et al. 1977, 1982; Elias et al. 2012; Hudek et al. 2016; Yan et al. 2017). In presence of arsenic, phosphate uptake is also modified. Under these conditions, arsenate competes for phosphate transporters and enters cells preferably through the non-specific Pit transporter, which fails to distinguish between phosphate and arsenate (Cleiss-Arnold et al. 2010). On the other hand, under arsenate stress, the Pst system increases its specific phosphate affinity more than 100-fold for some microorganisms (Guo et al. 2011) and may even exceed 4500-fold for extremophile microorganisms such as GFAJ-1 (Elias et al. 2012). Moreover, genes related to arsenic metabolism (e.g., *aio* and *arr*) are contiguous to phosphate metabolism genes (like *pst*) in arsenic genomic islands (Li et al. 2013), indicating that there could be functional and/or evolutionary associations to allow for the adaptation under different arsenic and phosphate concentration conditions (Muller et al. 2007; Li et al. 2013). Nonetheless, this potential relationship, if any, between *aio*/*arr* genes and *pst* genes is still unknown. Considering this proximity of genes related to arsenic metabolism and phosphate transport genes in the arsenic genomic islands (Li et al. 2013), we investigated how the presence and/or abundance of these genes varies at different chemical conditions mainly referred to the arsenic and phosphate concentration.

To perform this goal, we deliver a comparative metagenomic study using samples from four High Altitude Andean Lakes (HAAL): Tebenquiche and Brava in Chile (Atacama salt flat), Diamante and Socompa in Argentina (Argentine Puna region). These lakes harbor polyextremophilic Andean Microbial Ecosystems (AMEs) -such as biofilms, microbial mats, microbialites and endoevaporites, among others-that tolerate high UV solar radiation, low oxygen pressure, extreme temperature fluctuation and high concentrations of heavy metals and metalloids such as arsenic (Farías et al. 2013, 2014, 2017; Albarracín et al. 2015; Fernandez et al. 2016; Rasuk et al. 2016; Rascovan et al. 2016).

## Materials and Methods

### Sampling

Tebenquiche Lake and Brava Lake microbial mat samples were obtained in November 2012. Sample coordinates: Tebenquiche (S 23°08’18.5” W 068°14’49.9”, Fig 1B) and Brava mat (S 23°43’48.2” W 068°14’48.7”, Fig. 1C). For DNA analyses, triplicate cores (2 cm^2^ each) were taken to a depth of 3 cm of the mat and were pooled prior to homogenizing in order to obtain representative samples. Homogenates used for DNA extraction were stored at -20 °C in the dark and processed within a week (Farías et al. 2014; Fernandez et al. 2016).

**Fig. 1.**
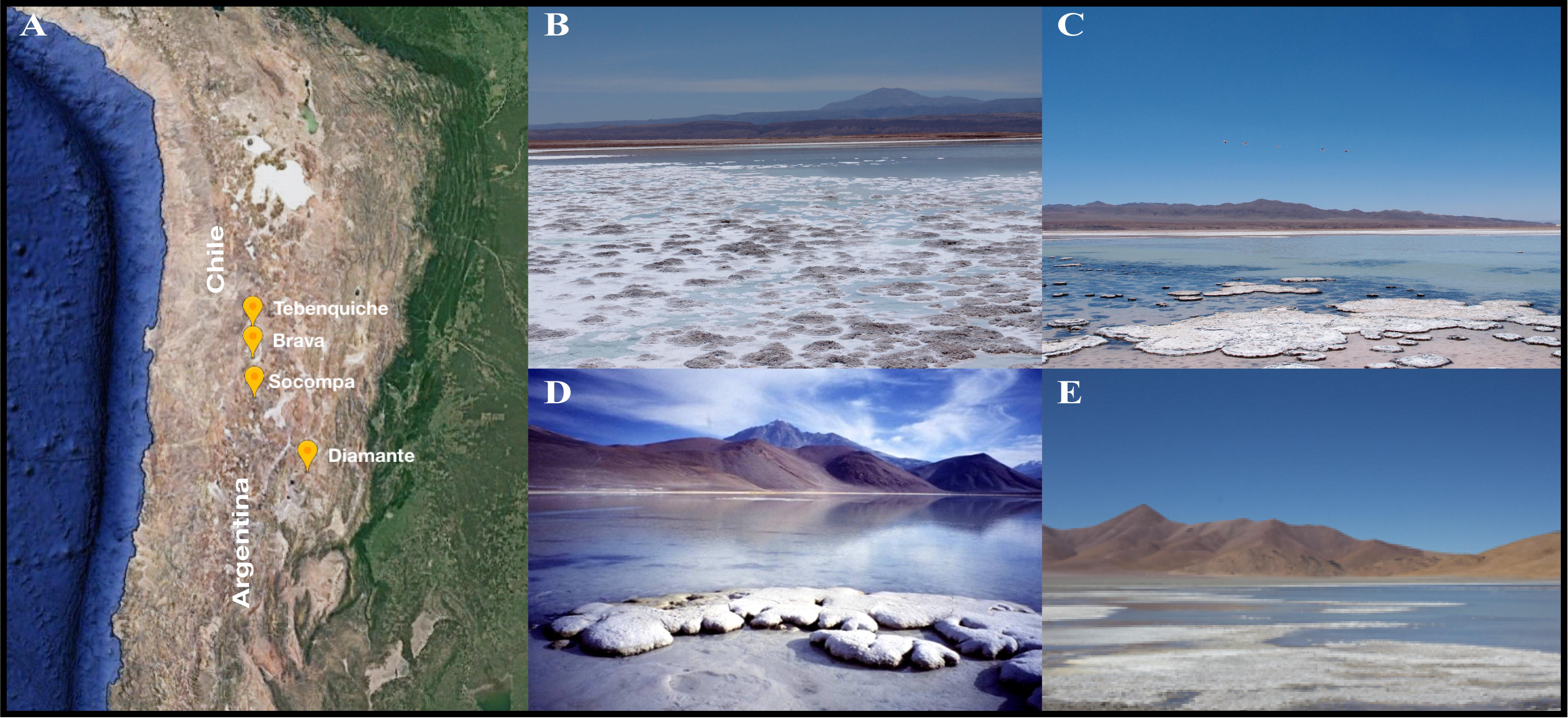
Location and photographs of the High-Altitude Andean Lakes under study. (A) Geographical location of the four lakes under study: Tebenquiche and Brava in Chile, Diamante and Socompa in Argentina. (B) Tebenquiche Lake; (C) Brava Lake; (D) Socompa Lake; (E) Diamante Lake. Photo credits to Farías ME and Saona LA.

Stromatolites can be found along the southern shore of Socompa Lake (coordinates: 24°32.168′S, 68°12.165′W, Fig. 1D). Samples were taken in February 2001, sampling methodology was already described by Farías (2013). These stromatolites are rounded domal structures that present a clear stratification that appears in vertical sections. Stromatolite samples were collected in sterile plastic bags and were fragmented into 2.5□cm-diameter cylinders that measured 5□cm deep from the top of the stromatolite. They were stored at -20 °C in the dark and processed within a week (Farías et al. 2013).

Biofilm samples were collected in November 2016 from Diamante Lake, located inside the Galan Volcano crater, located at 4589 masl (coordinates: 26 □00’09.71”S, 67 □02’34.91”W, Fig. 1E). Galan Volcano lies in the province of Catamarca, Argentina and reaches 5912 masl. Three independent samples were taken from Diamante Lake, and each was carried out by pooling together three representative samples. The samples were collected in centrifuge tubes, stored at -20 °C in the dark until they were processed (Rascovan et al. 2016).

For phosphate and arsenic content analysis, all water samples (1 L) were collected immediately over the sampling sites of the corresponding sediment systems and stored in acid-cleaned bottles on ice in the dark.

### Water physicochemical analysis

Tebenquiche and Brava water samples were stored in acid-cleaned bottles, on ice and in the dark until analyses in the laboratory were carried out within 48 h. Dissolved oxygen, salinity, conductivity, total P, NO−3, NO−2, dissolved Si, Ca2+, Mg2+, K+, SO2−4, and Na+, were measured according to the methodology described by Eaton et al. (2005). NH+4, orthophosphates, and Total Organic Nitrogen (TON) were analyzed using a Merck Nova 60 Spectro Photometer by following standard methods, as described by the American Public Health Association (1998).

For the Socompa Lake and Diamante Lake samples: nutrients, ions and general chemical analyses (arsenic and phosphate) were performed in a chemical IRAM-certified laboratory at Estación Experimental Obispo Colombres, Tucumán, Argentina (http://www.eeaoc.org.ar/).

### DNA extraction and sequencing

For all samples, total metagenomic DNA used for sequencing was extracted from a mixture of three replicate extractions using Power Biofilm DNA Isolation Kit (MO BIO Laboratories, Inc.) from 0.5g of collected biofilm/stromatolites/microbial mat which was processed according to manufacturer’s indications. From the biofilm of Diamante Lake, 3 independent extractions from 3 different replicates were made. In the case of stromatolites samples, first they were frozen and then were lyophilized and homogenized into a powder that was processed (Farías et al. 2013).

Total DNA was sheared by sonication with a target fragment size of ∼400 bp. DNA fragment distribution was confirmed in a Fragment Analyzer (Advanced Analytical) and libraries were constructed using the Illumina TruSeq DNA kit 2×300 bp as per manufacturer’s instructions. Libraries were multiplexed and run in an Illumina MiSeq instrument. Raw data for Diamante Lake’s red biofilm is available from NCBI’s SRA SRP136179 under BioProject PRJNA438526; for Tebenquiche and La Brava is available from ENA European Nucleotide Archive (ENA) under accession number: PRJEB25599; and for Socompa’s stromatolites is available from NCBI’s SRA SRP072938 under BioProject PRJNA317551.

### Metagenomic analyses

Metagenome raw data from each lake were filtered at PHRED > Q20 using Prinseq version 0.20.4 (Schmieder and Edwards 2011). Then, reads were assembled using 3.9.0 SPAdes version (Bankevich et al. 2012) with the --meta parameter to call the metaSPAdes module. The four assemblies were annotated using Prokka version 1.11 (Seemann 2014), adding the --metagenome parameter to improve gene prediction. Since many genes involved in arsenic and phosphate metabolism are known, we quantified their difference in abundance among the four metagenomes studied. Phosphate metabolism genes included in the analysis were: *ppk, ppx, pitA, phoB, phoR, pstS, pstC, pstA, pstB* and *phoU*; and the genes related to arsenic were: *acr3, aioA, aioB, arrA, arsR, arsD, arsA, arsB* and *arsC*. Relative abundances of these genes were obtained by mapping the reads against the contigs of each lake metagenome using bowtie2 (Langmead and Salzberg 2012). The raw counts of each gene were calculated with a htseq-count, a module of HTSeq (Anders et al. 2015). We used DESeq2 (Love et al. 2014) to normalize sequence coverage among the metagenomes and to perform inter-sample comparisons (negative binomial normalization; parametric Wald test at FDR < 0.01; Benjamini-Hochberg correction for multiple testing). Intra-sample comparisons for gene abundance analysis were performed using the FPKM metric. The taxonomic profile of the lakes was obtained by recruiting the small ribosomal sub unit gene sequence as implemented in metaxa2 (Bengtsson-Palme et al. 2015) by aligning the filtered reads against archaea, bacteria and eukaryote databases (parameter -t a,b,e of metaxa2). The resulting OTUs were analyzed in R using the phyloseq package (McMurdie and Holmes 2013).

## Results

### Characteristics of High-Altitude Andean Lakes

The HAAL studied here are characterized as shallow saline or thalassic lakes located in the Chilean Atacama Desert and in the Argentine Puna (Fig. 1A). All of them represent practically unexplored environments that have a set of unique extreme characteristics. Tebenquiche and Brava Lakes (Fig. 1B and 1C, respectively) are located in the lower region of the Atacama basin (Chile). Lake water comes from underground sources of tertiary and quaternary volcanic origin (Risacher and Alonso 1996) and they are characterized by a high concentration of lithium, boron and arsenic (Lara et al. 2012; Farías et al. 2014). Specifically, Tebenquiche Lake is one of the largest water bodies in the Atacama salt flat (Demergasso et al. 2008). Socompa Lake (Fig. 1D) is located at 3.800 meters above sea level in a basin at the base of the active Socompa Volcano (Argentina). As with the rest of the lakes, arsenic concentration is very high (Farías et al. 2013). Finally, Diamante Lake (Argentina) (Fig. 1E) is located at 4.570 meters above sea level within the crater of the Galan Volcano, near a hydrothermal spring effluent. Apart from sharing all the HAAL characteristics, its arsenic concentration is extremely high (Fig. 2), more than 10 times the concentration of Mono Lake, a habitat that features high arsenic content (Wolfe-Simon et al. 2011; Rascovan et al. 2016).

**Fig. 2.**
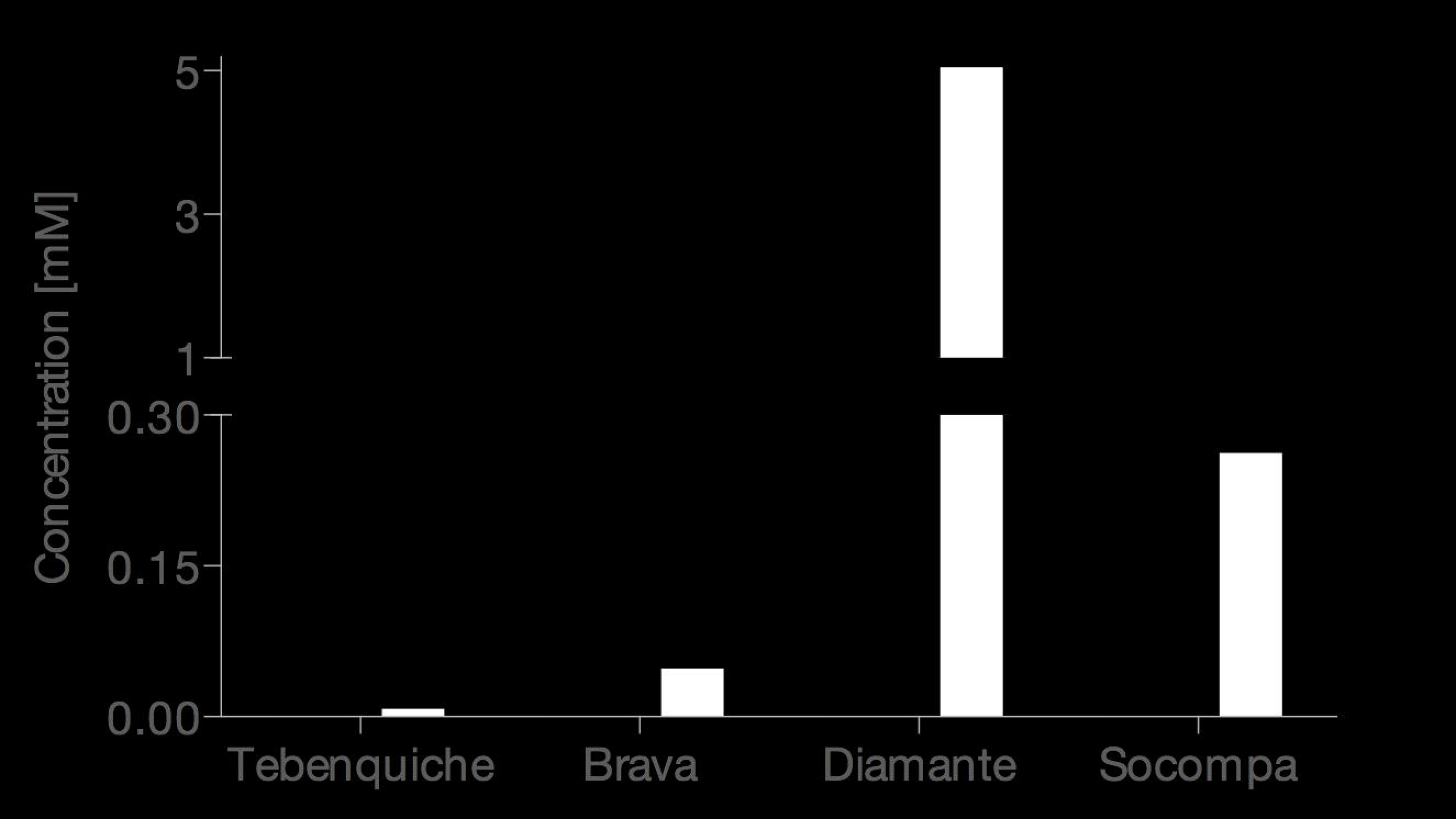
Chemical analysis of samples used for metagenomic studies. Phosphate and arsenic concentration of the samples obtained from Tebenquiche, Brava, Diamante and Socompa lakes were quantified. Black bars represent arsenic concentration and white bars phosphate concentration.

In order to determine arsenic and phosphate concentrations in the four Andean lakes, a physicochemical analysis was performed which revealed that Tebenquiche, Brava and Socompa (TBS) lakes have low concentrations of both phosphate and arsenic in comparison to Diamante Lake. Tebenquiche showed the lowest concentrations with 0.011 mM and 0.053 mM of phosphate and arsenic, respectively. On the other hand, the concentration of phosphate and arsenic in Diamante Lake was 5.100 and 2.340 mM, respectively (Fig. 2), which shows that, compared to Tebenquiche, arsenic concentration is 40-fold higher.

### Taxonomic compositions of metagenomes

We found significant taxonomic similarities between TBS lakes, whereas Diamante Lake displayed differences regarding the dominant phyla, mainly due to the high abundance of the Euryarchaeota phyla. This observation agrees with previous reports pertaining Diamante Lake (Rascovan et al. 2016). On the other hand, Fig. 3B shows the most abundant taxonomic groups per sample classified at the family level, revealing large differences among all the lakes with a predominant Halobacteriaceae family in Diamante Lake. While TBS are dominated by bacteria, Diamante is dominated by archaea, where the most abundant phylum is Euryarchaeota. Indeed, this phylum represents 94% of the total microbial community (Rascovan et al. 2016). In Diamante, the only phylum belonging to the bacteria domain is Proteobacteria represented in more than 5%. At the family level, the taxonomic similarity between TBS is not as evident as it is at the phylum level with the 5 most abundant families that represent 49.7%, 43.5% and 39.7%, respectively. Spirochaetaceae is the unique family shared by Tebenquiche and Socompa (Fig. 3B). There are also no major taxonomic similarities between the studied lakes when examining the most abundant families (Fig. S1). Alpha diversity consistently showed that Diamante is less diverse compared to the other lakes (Fig. 3C; using 4 diversity metrics), which in turn concurs with the taxonomic profile at the Phylum and Family levels (Fig. 3A and 3B). Altogether, this suggests that Diamante Lake imposes harsh conditions for microbial life to thrive.

**Fig. 3.**
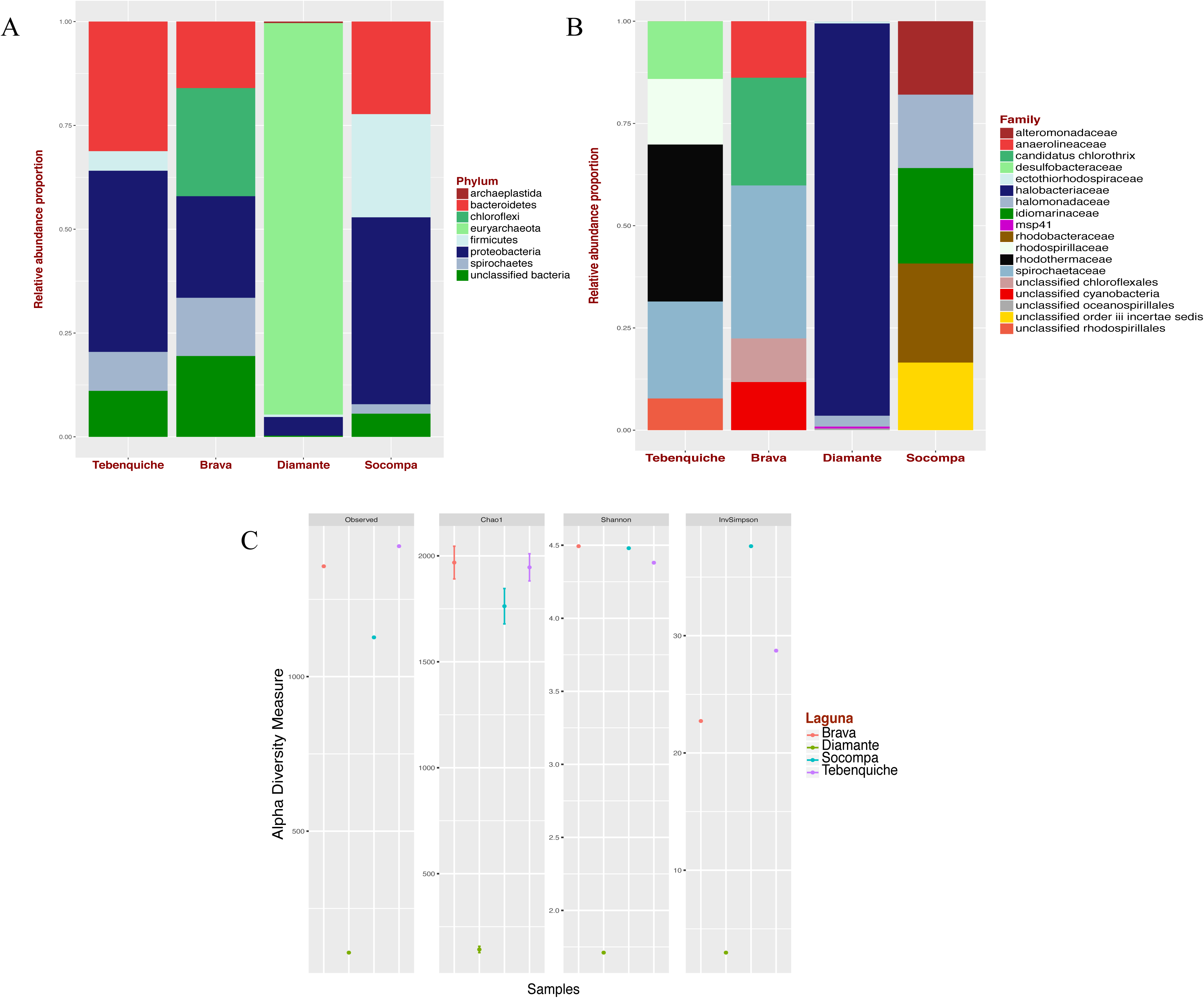
Comparison of microbial diversity in Tebenquiche, Brava, Diamante, and Socompa lakes. (A) Stacked bar chart representing relative abundance of dominant phyla in the four lakes. (B) Stacked bar chart representing relative abundance at the family level. (C) Alpha diversity with four measures: Observed, Chao1, Shannon and InvSimpson.

### Phosphate metabolism genes

*PhoR*/*phoB* genes belong to the phosphate (Pho) regulon, and are implicated in bacterial/archaeal phosphate regulation (Wanner and Chang 1987; Santos-Beneit 2015). Through *phoR* and *phoB* genes, the Pho regulon is involved in phosphate concentration sensing on the periplasm (Lamarche et al. 2008). TBS lakes have much less phosphate than Diamante Lake (Fig. 2) and exhibit higher *phoR* and *phoB* gene abundance. The *phoR* gene is 7.26, 16.57 and 3.46 times less abundant in Diamante than in TBS, respectively. Likewise, the abundance values for the *phoB* gene are 13.54, 25.82 and 21.26. On the other hand, another key molecular player in phosphate metabolism is the PhoU enzyme –a metal binding protein– that is involved in the Pst phosphate transport system and belongs to the *pstSCAB-phoU* operon (Steed and Wanner 1993; Lamarche et al. 2008; McCleary 2017). Unlike *phoR* and *phoB*, the *phoU* gene is 280.15, 689.69 and 146.02 times more abundant in Diamante than in TBS lakes, respectively (Fig. 4), suggesting that Diamante might favor the transcription of genes related to specific phosphate transporters (Pst) by means of PhoU, whereas, the synthesis of these transporters in the other lakes might be very well regulated by PhoB and PhoR.

**Fig. 4.**
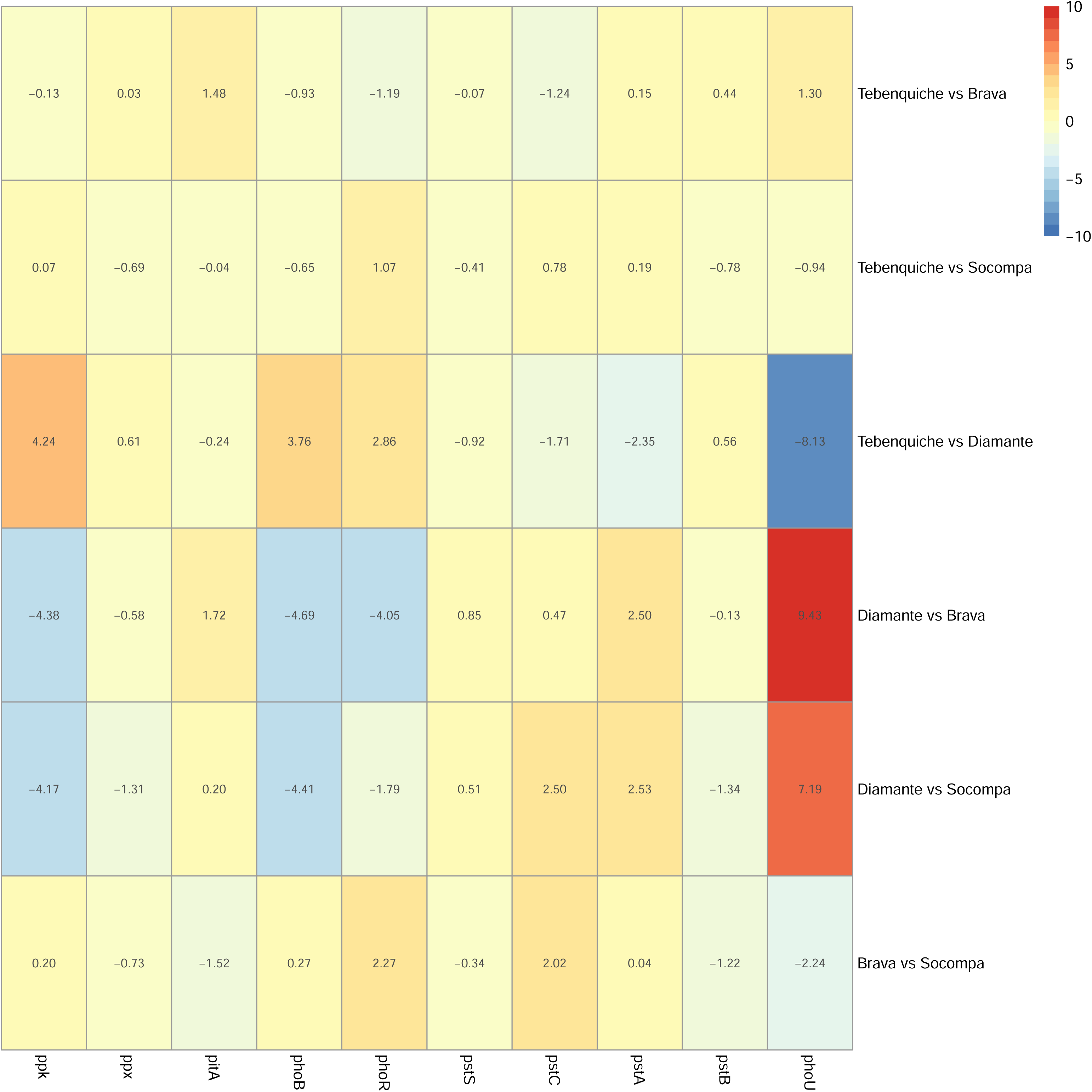
Differential abundance analysis of genes associated with phosphate metabolism. The differential abundance of the *ppk, ppx, pitA, phoB, phoR, pstS, pstC, pstA, pstB* and *phoU* genes in all the lakes studied was calculated. The results were plotted by heatmap when comparing one lake against another. The biggest differences are in the Diamante Lake.

The genes of the *pst* operon (*pstS, pstC, pstA* and *pstB*) encode for a high-affinity ABC type phosphate transporter. Specifically, the *pstS* gene codes for a periplasmic phosphate-binding protein, whose function is to bind phosphate from the external medium and transfer it to another protein complex (PstCAB) that will enter the cell cytoplasm (Blus-Kadosh et al. 2013). Our results, unexpectedly, show that the *pstS* gene is 1.90, 1.81 and 1.43 times more abundant in Diamante than in TBS. The *pstC* and *pstA* genes coding for inner-membrane channel proteins for Pst transport (Lamarche et al. 2008) are more abundant in Diamante than in the other lakes, with the exception of Brava Lake that has 1.49 times more *pstC* than Diamante. The *pstC* gene is 3.27 and 5.66 times more abundant in Diamante than in Tebenquiche and Socompa, respectively; whereas *pstA* is 5.10, 5.66 and 5.78 times more abundant in Diamante than in TBS, respectively (Fig. 4).

Regarding the Pit system –the low-affinity phosphate transport–, the *pitA* gene is 1.18, 3.29 and 1.15 times more abundant in Diamante Lake than in TBS, respectively (Fig. 4). The intra-sample comparison reveals a very low relevance of the Pit transporter in all lakes, moreover, the abundance of the *pitA* gene is the lowest among the phosphate metabolism genes evaluated (Fig. S2), which suggests that phosphate uptake would take place mostly by means of the Pst system.

Finally, as to polyphosphate metabolism genes –*ppk* and *ppx*–, both show very similar abundance in TBS lakes, while at Diamante abundance is very low. Specifically, the *ppk* gene is 18.89, 20.83 and 18.01 times less abundant in Diamante than in TBS lakes, respectively; in the same order, the values for the *ppx* gene are: 1.53, 1.50 and 2.48.

### Arsenic resistance and metabolism genes

The analysis of *ars* operon genes reveals great differences between the four metagenomes studied. The *arsR* gene has a similar abundance in Diamante, Tebenquiche and Brava lakes, while in Socompa it is 2.33, 1.34 and 2.89 times more abundant, respectively. The *arsD* and *arsA* genes are very well distributed in Diamante compared to the other lakes; *arsD* is 4.86, 17.39 and 1.56 times more abundant in Diamante than in Tebenquiche, Brava and Socompa, while in the same order, the values for the *arsA* gene are: 7.68, 9.72 and 6.50. The *arsC* gene is significantly more abundant in the lakes where arsenic concentration is low, as it is 24.30, 34.30 and 15.78 times more abundant in Tebenquiche, Brava and Socompa, respectively, than in Diamante. The arsenate reductase enzyme, ArsC, is responsible for reducing As(V) into As(III) which is subsequently expelled from the cell by the ArsB efflux pump. Though the *arsB* gene was not found in Diamante Lake, intra-sample comparisons made using the FPKM method show that the *arsB* gene is not abundant in the other three lakes either (Fig. S3).

In addition to the ArsB efflux pump, there is another arsenic efflux pump widely distributed among microorganisms: the Acr3. The *acr3* gene is 3.27, 3.21 and 3.21 times more abundant in Diamante than in Tebenquiche, Brava and Socompa, respectively (Fig. 5). Meanwhile, the abundance of the *acr3* gene is equivalent among Tebenquiche, Brava and Socompa, and its abundance in each individually is much higher than the abundance of the *arsB* gene (Fig. S3). This suggests that the expulsion of arsenic from the cytoplasm might be carried out mainly by the Acr3 pump, which is common to the 4 lakes.

**Fig. 5.**
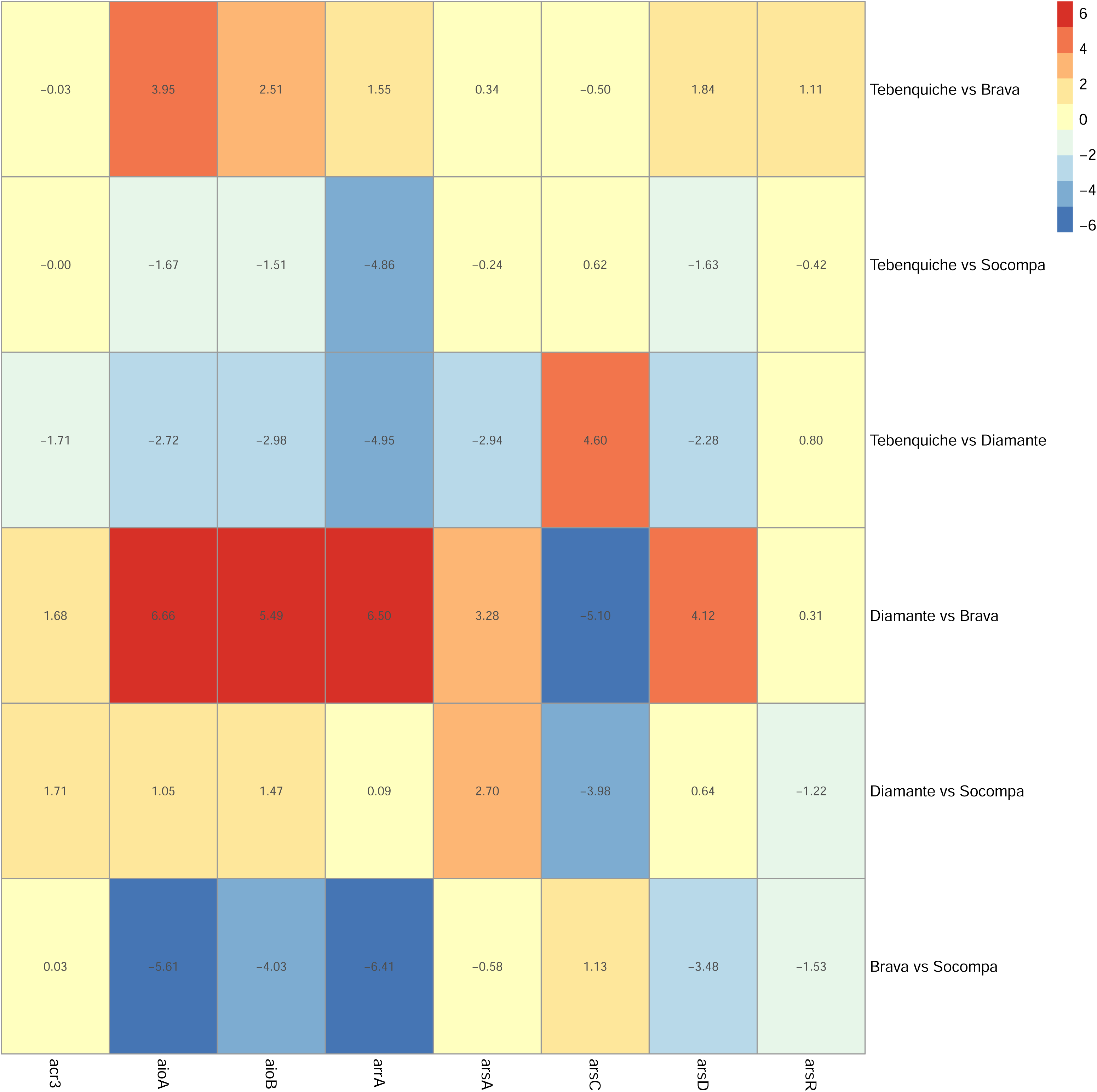
Differential abundance analysis of genes associated with arsenic metabolism. The differential abundance of the *acr3, aioA, aioB, arrA, arsA, arsC, arsD* and *arsR* genes in all the lakes studied was calculated. The results were plotted by heatmap when comparing one lake against another. The biggest differences are in the Diamante Lake.

Finally, the genes corresponding to arsenic respiration –*aioAB* and *arrA*– were analyzed (Rascovan et al. 2016). The *aioA* and *aioB* genes that encode an arsenite oxidase have a much greater relative abundance in Diamante Lake than in the others. Diamante and Brava show the largest difference, where *aioA* and *aioB* are 101.15 and 44.95 times more abundant in Diamante, respectively. On the other hand, the *arrA* gene coding for arsenate reductase presents a great difference as it is much more abundant in Diamante Lake and *arrA* is 30.91 and 90.51 times more abundant in there than in Tebenquiche and Brava. For its part, Socompa does not show significant differences with Diamante in terms of the *arrA* gene (Fig. 5).

## Discussion

Our work characterizes the microbial communities and analyzes the abundance of genes related to arsenic and phosphate metabolisms found in four salt lakes distributed along the Argentine Puna and the Atacama Desert in Chile. The results reveal that the greatest differences in gene abundance appear at Diamante Lake in contrast with TBS lakes (Fig. 4 and Fig. 5). These differences are also reflected in the distribution of species and biodiversity (Fig. 3); While archaea comprises 94% of Diamante (Rascovan et al. 2016), in the other lakes the main species are bacteria (Farías et al. 2013; Fernandez et al. 2016; Rasuk et al. 2016), and have a much higher biodiversity than Diamante (Fig. 3D). Finally, in terms of phosphate and arsenic concentration, physicochemical conditions also evidence considerable differences between Diamante and the other lakes (Fig 2).

The *aioA*/*B* genes encode proteins that use As(III) as an electron donor, oxidizing it to As(V). Because these enzymes have been described as transmembrane proteins (Andres and Bertin 2016), this process occurs within the extracellular environment in the presence of arsenic. As(V), on the other hand, is a substrate for another family of transmembrane enzymes, the ArrA/B, which use As(V) as the final acceptor of the electron transport chain and reduce it to As(III). The theorization of this process, has shown that cells can *breathe* arsenic and thereby obtain energy from its oxidation and reduction (Rascovan et al. 2016; Andres and Bertin 2016). The *aio* and *arr* genes are very well distributed in Diamante Lake and their abundance is much higher than in the other lakes analyzed (those with the lowest arsenic concentration) (Fig. 2). In summary, the higher the arsenic concentration, the higher *aio* and *arr* gene abundance.

Regarding the *arsRDABC* operon, both *arsA* and *arsD* genes are highly abundant in Diamante compared to the other lakes, yet the same does not apply to *arsR, arsB* and *arsC* genes. The *arsRBC* operon confers basal tolerance to arsenic, mainly by means of the arsenate reductase ArsC, which is in charge of the electrochemical transformation of As(V) to As(III), subsequently expelled from the cell by the ArsB efflux pump (Dey and Rosen 1995; Lin et al. 2006; Andres and Bertin 2016). These genes belong to the *arsRBC* operon and their abundance in Diamante is very low, particularly *arsC*. As a matter of fact, *arsB* is simply not present, even when mapping the reads against a Hidden Markov Model (HMM) of the *arsB* gene, no matches are found. Although *arsB* did not turn out to be a particularly abundant gene in TBS lakes, it is very likely that it is not present in Diamante. The Diamante metagenome was sequenced in triplicate from three different samples to minimize the risk of not finding a gene due to lack of sequencing (BioSamples accessions SAMN08719551, SAMN08719552 and SAMN08719553).

Unexpectedly, arsenic resistance genes from *arsRBC* operon were (in the case of *arsC*) up to 35 time more abundant in lakes with low arsenic concentration, while in Diamante, where arsenic concentration is up to 40 times greater than in the other lakes, only genes related to oxidation and reduction of extracellular arsenic are more abundant. Previous works have demonstrated that *arsC* activity is modified in the presence of phosphate (Zhang et al. 2014, 2017); it may be the case that despite high arsenic concentration, there is low arsenic uptake by the cells, rendering an arsenate reductase (ArsC) not vital for survival. In addition, the expulsion of arsenic from the cell -in all lakes-might not be performed by ArsB but by Acr3, a pump that does not belong to the *arsRBC* operon. An intra-sample analysis shows that the *acr3* gene is very well distributed in all lakes, especially in Diamante (Fig. S3). Previous findings have shown *acr3*-gene abundance in *Exiguobacterium* strain genomes (Ordoñez et al. 2015; Castro-Severyn et al. 2017), nevertheless, further analyses are necessary to understand the relationship between the *arsB* and *acr3* genes, as well as their abundance according to the physicochemical conditions of different environments.

In the case of phosphate transport systems, both the Pst and Pit, are distributed in the four metagenomes studied. However, while the Pit system does not show remarkable differences among the 4 lakes, the Pst system is high abundant in Diamante Lake in comparison with TBS lakes (Fig. 4). The Pst is a high affinity and a low speed transport system, while the Pit transport system is characterized as a constitutive shock-resistant system which does not depend on membrane-bound proteins like the Pst (Lamarche et al. 2008; McCleary 2017). Previous studies have shown that the Pst system is activated in conditions of phosphate deprivation (Sprague et al. 1975; Richards and Vanderpool 2012; Elias et al. 2012; Reistetter et al. 2013), however in presence of high phosphate concentration it remains active playing a key role (Guo et al. 2011; Hudek et al. 2016). Based on this, apparently, the Pst system would be key to the phosphate uptake at low and high phosphate concentrations (Bergwitz and Jüppner 2011). This is consistent with our results, why in the Diamante Lake, where the phosphate concentration is excessively high, is there a greater abundance of Pst genes compared to the TBS lakes?

On the other hand, studies with *Halomonas* GFAJ-1 have discovered the presence of two *pst* operon, where the PstS phosphate binding protein of one of them selects phosphate at least 500-fold over arsenate (Elias et al. 2012), whereas the PstS belonging to the Pst2 operon shows 4,500 times more affinity (Wolfe-Simon et al. 2011; Yan et al. 2017). As mentioned by Schoepp-Cothenet (2011), it seems that the *Halomonas* GFAJ-1 has evolved to incorporate enough small amounts of phosphate to survive while intoxicating with arsenic (Schoepp-Cothenet et al. 2011); this is why GFAJ-1 would be able to grow with trace phosphate concentrations (3 μM) even in high arsenic concentration. Our results support that hypothesis. In Diamante Lake where arsenic concentration is extremely high (2.34 mM) and despite the high phosphate concentration, the Pst system is more abundant in comparison with the other lakes. The intra-sample analysis in Diamante Lake reveals the *pstSCAB-phoU* operon genes are the most relevant genes belonging to phosphate metabolism, and their abundances are even greater than the *dnaG* housekeeping gene (Fig. S2).

Under arsenic stress, cells need to have more specific phosphate uptake systems (Li et al. 2013), besides, through the transcriptional profile of *H. arsenicoxydans* it was observed that under conditions of As(V) exposition, phosphate uptake occurs preferentially through the specific Pst system rather than the Pit general transport, in order to reduce the entry of As(V) (Cleiss-Arnold et al. 2010). This phenomenon could mean that evolutionarily the microbial life in Diamante Lake had to select specific phosphate transporter system to avoid arsenic uptake. By means of *in silico* analyses of bacterial genomes, arsenic islands have been discovered where *pst* genes are next to *aio* genes (Li et al. 2013; Chen et al. 2015). The relation of the *aio* genes with the *pst* operon around the arsenic islands is not yet elucidated (Lebrun et al. 2003; Kim and Rensing 2012), however, as other authors have suggested, it is possible that through the arsenic islands, arsenic induces production of Pst transporters that allows the cells to obtain the phosphate they need in arsenic laden hypersaline lakes (Willsky and Malamy 1980a; Cleiss-Arnold et al. 2010; Li et al. 2013).

Regarding the *pst* operon regulator, Diamante lake shows a huge abundance of *phoU* gene and a very low presence of *phoR*/*phoB* genes (Fig. 4). This is consistent with expectations since *phoR*/*phoB* genes are expressed under conditions of low phosphate concentration (Lamarche et al. 2008; Chen et al. 2015; McCleary 2017), while *phoU* are expressed under conditions of high phosphate concentration (Steed and Wanner 1993; Lamarche et al. 2008). In addition, PhoU is essential to avoid an uncontrolled uptake of Pi that could become toxic to the cells (Surin et al. 1986; Santos-Beneit 2015), which is what could be happening in Diamante Lake. However, the whole functions of PhoU are not elucidated yet, even less its relationship with arsenic metabolism. On the other hand, the PhoB protein is essential for polyP accumulation, in fact, *E. coli* mutant strains lacking *phoB* do not have the ability to accumulate intracellular polyP (Rao et al. 1998). The polyP metabolism genes (*ppk* and *ppk*) are not well distributed in Diamante Lake (Fig. S2), instead both genes are very abundant in TBS Lakes (Fig. 4).

High expression of phosphate metabolism-related genes (Pst system and Pho regulon mainly) influenced by presence of arsenic in microorganisms has been previously verified (Cleiss-Arnold et al. 2010; Kang et al. 2012a; Li et al. 2013). In the same way, the arsenic metabolism (*arsC* mainly) affected by the phosphate concentration has been also documented (Markley and Herbert 2010; Slaughter et al. 2012; Kang et al. 2012b; Wang et al. 2013; Zhang et al. 2014). In accordance with this, our results show that in the face of high phosphate and arsenic concentration: (i) the *aio* and *arr* genes are very abundant suggesting that obtaining energy from arsenic substrates would be a well distributed process, (ii) high specificity of phosphate transporters (Pst) would possibly restrict arsenic uptake, (iii) *phoU* gene is extremely abundant, this would act by regulating phosphate homeostasis avoiding high phosphate uptake (iv) *arsC* gene involving in the reduction of arsenate is not abundant suggesting that arsenic uptake would be restricted in this conditions. On the other hand, in face of low phosphate concentration (and more arsenic than phosphate), our results show: (i) *aio* and *arr* genes are not abundant, (ii) *phoB*/*phoR* genes involving in the activation of Pst system in phosphate starvation are very abundant and finally, (iii) *arsC* is highly abundant, maybe because in these conditions the arsenic uptake is higher.

These results expand our understanding of how the arsenic and phosphate metabolism would be intricately co-regulated and the way that the environmental factors would influence on the microbial metabolism through genes selection. With this manuscript, we attempt to contribute to the current discussion about As and P metabolism, specifically in polyextremophile conditions.

## Acknowledgements

ECN was funded by “CONICYT-FONDECYT de iniciación en la investigación 11160905”. ECN would like to thank George Washington University’s high-performance computing facility, Colonial One, for providing data storage, support, and computing power for metagenomic analyses (colonialone.gwu.edu/).

## Author contributions statement

LAS contributed with the principal idea of this work, participated in the study design, performed data analysis, interpreted data and wrote the paper. SVD contributed with design methodology for data analysis, performed data analysis, and contributed with article writing. DK contributed with the work proposal. MC obtained funding for the original project idea and performed the physicochemical analysis. CM contributed with sequencing of some metagenomes. ECN participated in the study design, choice of methodology and data analysis, sequenced some of the metagenomes, and helped write the manuscript. MEF obtained funding for the original project idea, contributed with the work proposal, with the sampling and sequencing of the metagenomes. All authors read and approved this manuscript.

