## Supplementary figures and images for "Analysis of co-regulated abundance of genes associated with arsenic and phosphate metabolism in Andean Microbial Ecosystems"

### Supplemental Figure 1

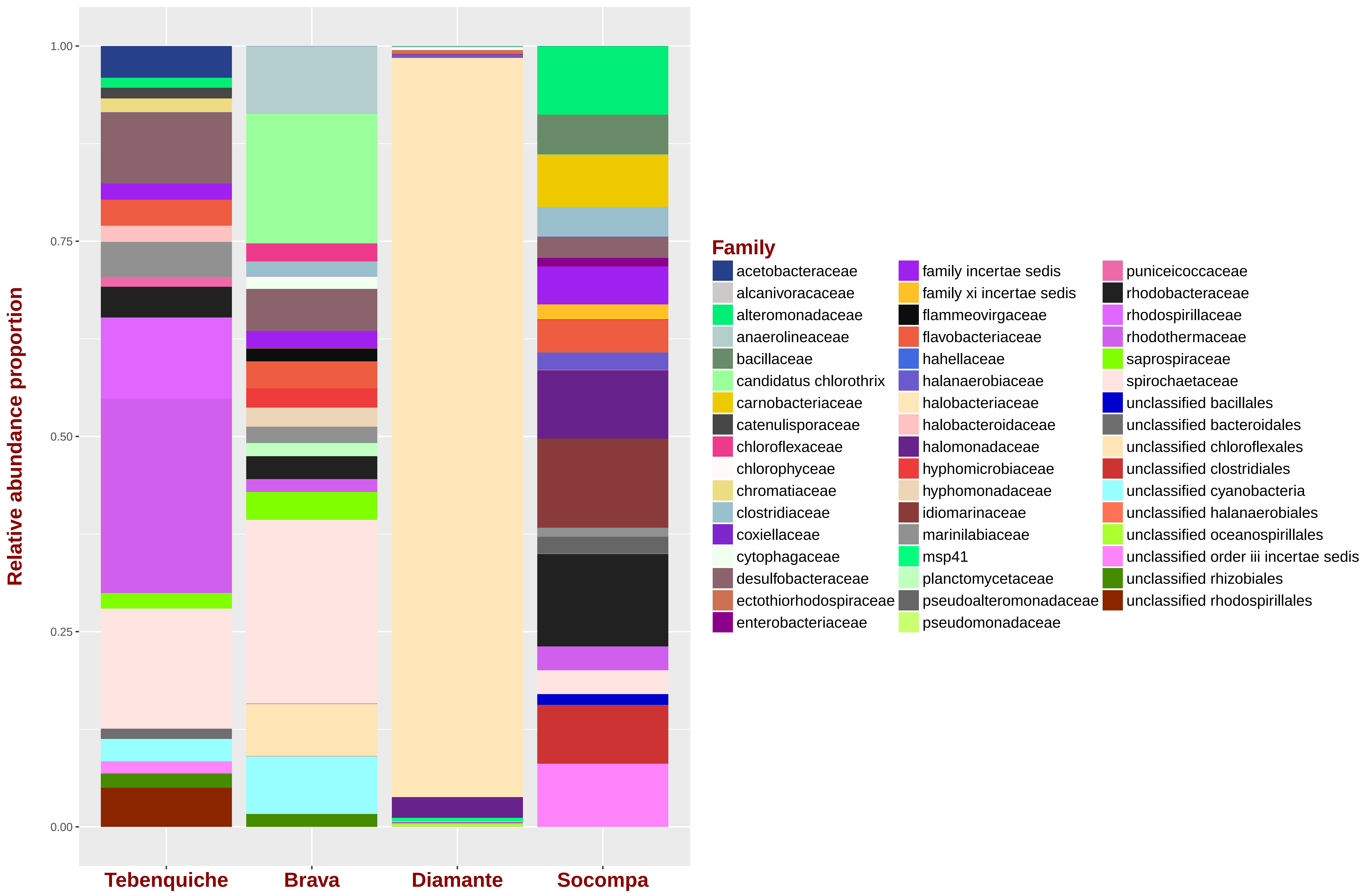

### Supplemental Figure 2

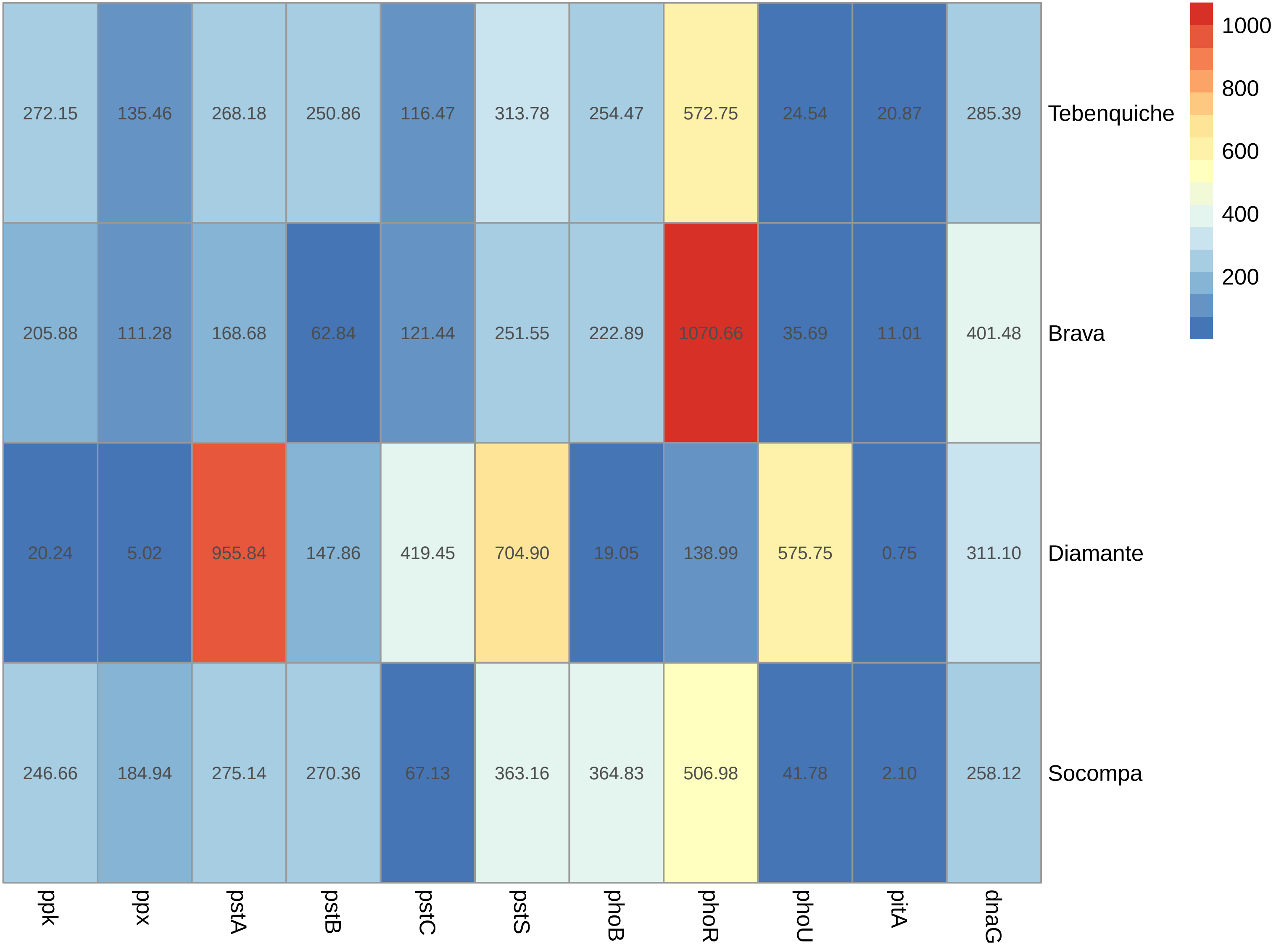

### Supplemental Figure 3

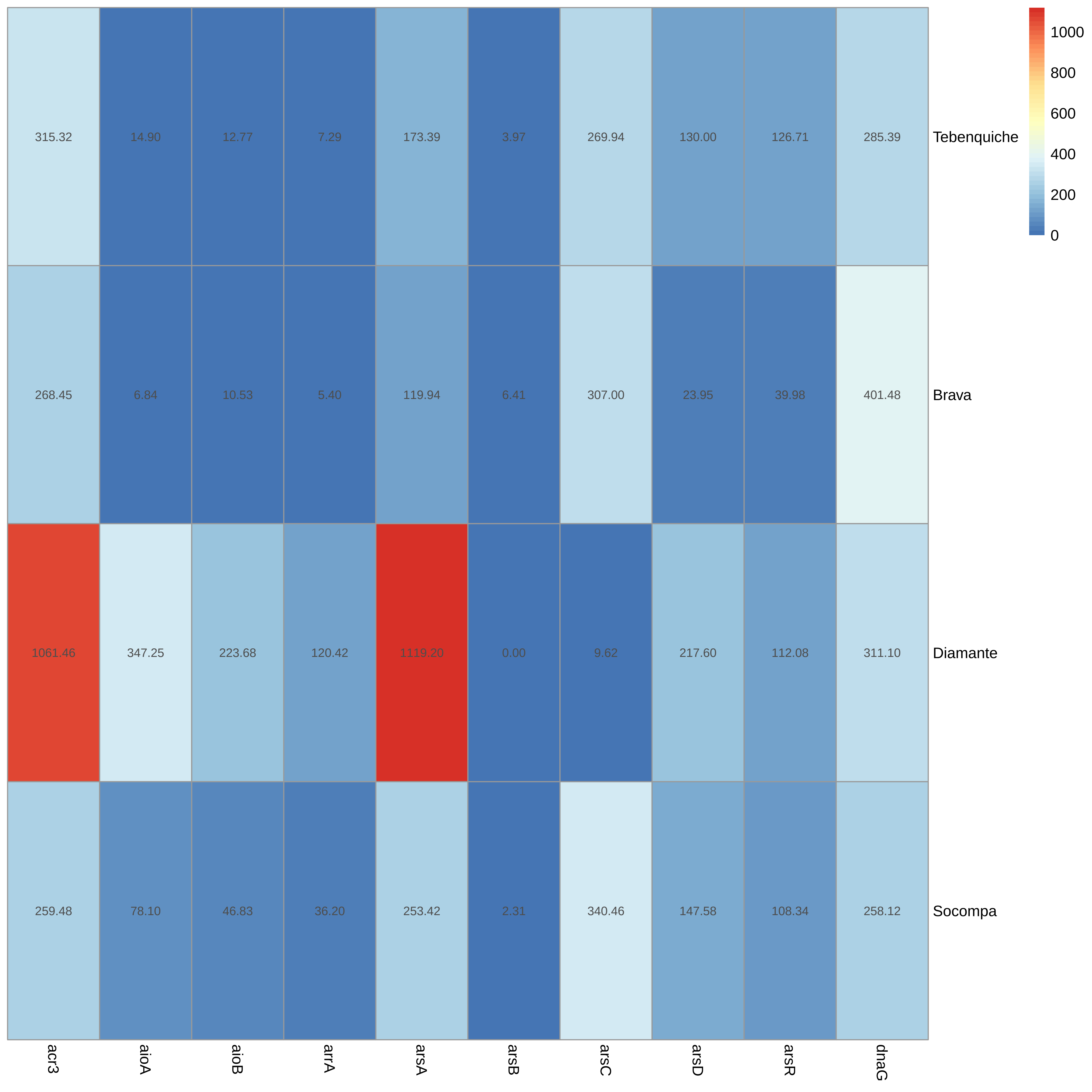
